# Root meristem shaping via brassinosteroid-controlled cell geometry

**DOI:** 10.1101/2021.04.01.438011

**Authors:** Y. Fridman, S. Strauss, G. Horev, M. Ackerman-Lavert, A Reiner Benaim, B Lane, R.S. Smith, S. Savaldi-Goldstein

## Abstract

Growth extent and direction determine cell and whole-organ architecture. How they are spatiotemporally modulated to control size and shape? Here we tackled this question by studying the effect of brassinosteroid (BR) signaling on the structure of the root meristem. Quantification of the 3D geometry of thousands of individual meristematic cells across different tissue types showed that modulation of BR signaling yields distinct changes in growth rate and anisotropy, which affects the time cells spend in the meristem and has a strong impact on final root form. By contrast, the hormone effect on cell volume was minor, establishing cell volume as invariant to the effect of BR. Thus, BR has highest effect on cell shape and growth anisotropy, regulating overall radial growth of the meristem, while maintaining a coherent distribution of cell sizes. Moving from single-cell quantification to the whole organ, we developed a computational model of radial growth that demonstrates how differential growth regulation by BR between the inner and outer tissues shapes the meristem. The model explains the unintuitive outcomes of tissue-specific perturbation of BR signaling and suggests that the inner and outer tissues have independent but coordinated roles in growth regulation.

## Introduction

Plant morphogenesis is determined by the rate of growth (cell expansion and cell division) and its directionality (anisotropy) ^1^. Growth rates are governed by hormonal signaling, the decoding of which depends on hormone levels and on the tissue and cell type in which it occurs. How hormonal signaling coordinates anisotropy remains unclear. The primary root meristem is composed of concentric tissue files surrounding the inner-most stele cells (Fig. 1). In the longitudinal axis, stem cell daughters undergo a series of anticlinal divisions in their corresponding tissue file before they begin to rapidly elongate in the elongation zone ^2^. The root meristem also expands in width by a series of radial (periclinal) divisions that increase the number of procambium cells in the stele, and by tangential divisions that add additional cell files to select tissues, involving interwoven transcriptional factors and hormonal signals ^3, 4^.

**Fig. 1.**
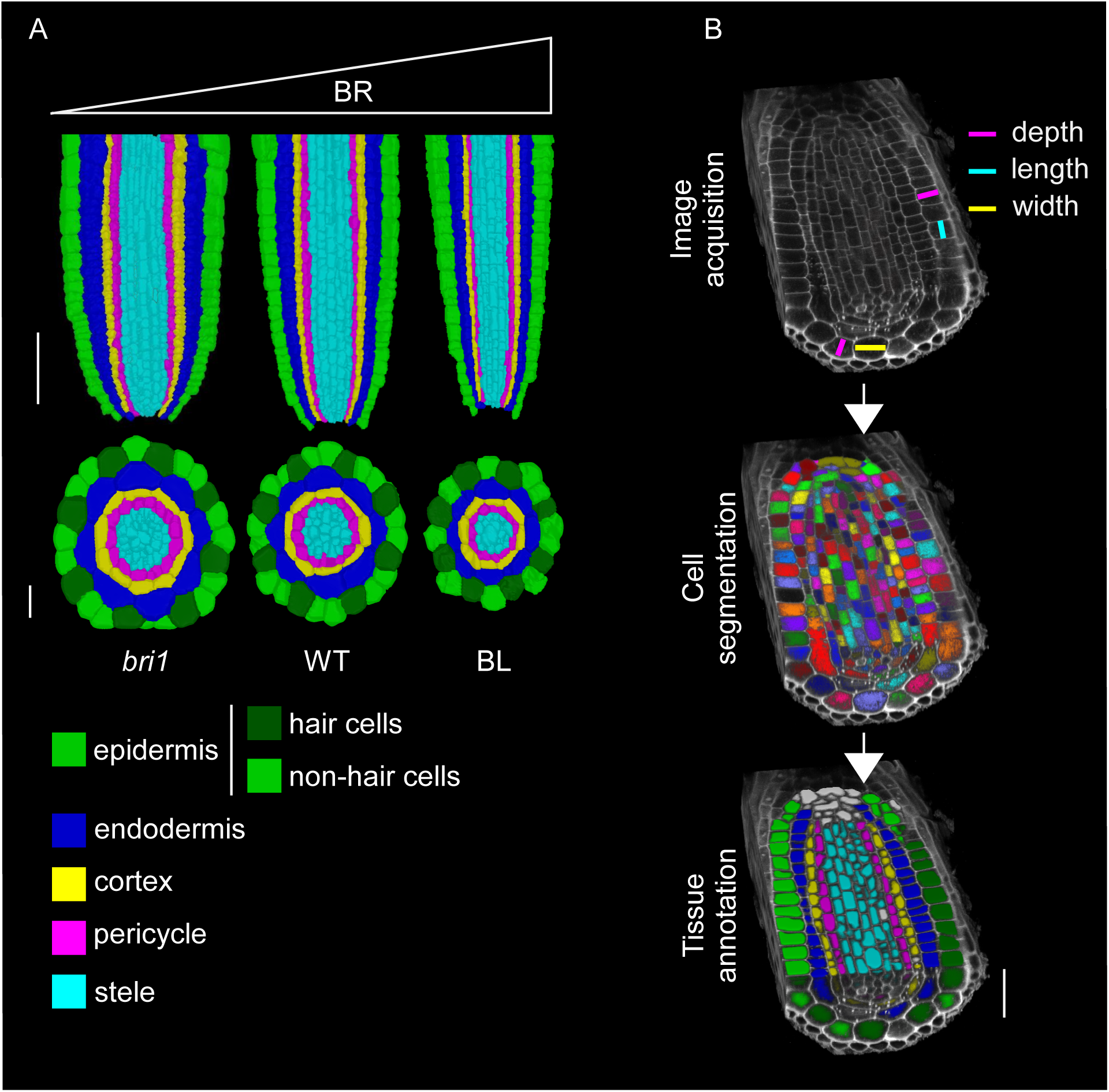
Root meristem morphology and segmentation scheme (A) Confocal images presented as longitudinal (upper panels) and radial (lower panels) cross-sections of root tip of a 7-day-old seedling, showing morphological differences between *bri1,* WT and WT grown in the presence of BL for 4 days. Cells underwent membrane-based segmentation and classified into different tissues, as shown by different pseudo-colors. Note the decreasing root diameter with increasing BR signaling. longitudinal scale bar = 50 μm, radial scale bar = 20 μm. (B) Summary of key steps to obtain tissue-specific 3D geometric parameters. Marked are volumetric cell geometries and their positioning in root cells. Scale bar = 20 μm.

As cells grow in a tissue context, they are subjected to mechanical feedbacks ^5–7^, which control whole-organ shape. Kinematics and additional quantifications of root growth parameters have been used to assess relative changes in growth rates among genotypes and treatments ^8^. However, the analysis of the effect of hormones on three-dimensional (3D) growth on a cellular scale is just emerging ^9^ and data on the accompanying changes in cell volume and growth anisotropy are lacking.

Brassinosteroid (BR) signaling regulates cell length and meristematic cell counts in the root in both the longitudinal and radial axes ^10, 11^. The signaling is initiated upon binding of the hormone to its cell surface receptor BRI1 through a regulatory sequence involving inactivation of the GSK3 kinase BIN2, which plays a major inhibitory role by phosphorylating and thereby inhibiting the activity of key downstream transcription factors belonging to the BES1/BZR1 family ^12^. High BR levels limit the number of dividing cells in the longitudinal and radial axes, promote early exit from the meristem and increase cell length ^13–15^. The *bri1* mutant has a short meristem ^13, 16^, with an increased number of cells in the radial axis ^14, 17^. Morphologically, these meristems have longer cells, arranged within a narrow structure (high BR), or have shorter cells arranged within a wider structure (low BR). However, BR signaling at the cellular scale has non-intuitive effect on the whole-root meristem structure ^10, 18^. Specifically, limiting BRI1 expression to the outer tissues promotes meristem length and restricts the meristem width while limiting BRI1 to the stele had the opposite effect ^16, 18^ resulting in a meristem structure, that is wider than that of *bri1* ^16, 17^. Similarly, inhibition of BR signaling via expression of the dominant-active version of BIN2 in the outer and the inner tissues only, yielded wider and narrower meristems than wild type (WT), respectively ^19^, highlighting the essential role of the outer tissues in restricting radial growth. A recent model proposes that BR regulation of BR biosynthesis genes in the inner stele tissue modulates BR levels that are perceived in the outer tissues, thus providing one mode of inter-tissue coordination. In addition, the levels of local BR production in the meristem are important for optimal root growth ^15, 20^ and BR intermediates appear to move within the meristem ^20^. However, data on how BR signaling controls geometry on a cellular scale and how it integrates to the whole-organ scale, as with radial growth of the root meristem, is lacking.

A better understanding of morphogenesis and the regulatory signaling involved requires precise single-cell tools that quantify growth parameters in 3D at the cell scale ^21–26^. However, their application is still scarce, and quantitative analysis of growth rates in 3D (i.e. 4D) is rarely performed, mainly due to difficulties of segmenting microscopy images. Here, we quantified the geometry of meristematic cells in Arabidopsis roots with adequate, low and high BR signaling, using 3D and 4D analyses. We then integrated experimental data in a computational model of radial meristem growth, and revealed how BR shapes the meristem at the cell level and that tissue specific constraints, modulated by BR, yield a coherent morphological output.

## Results

### BR modifies meristematic cell shape but not cell volume

MorphoGraphX ^27^ was deployed to precisely quantify the 3D geometry of meristematic cells in WT and BR-perturbed roots, and to compare the length, width, depth, surface area and volume of various tissues ^27^ (Fig. 1). Virtually all cells of the root meristem of independent WT and *bri1* roots and of WT roots treated with the BR brassinolide (BL), hereafter referred to as “treatments”, were segmented (with the exception of lateral root cap and stele cells inner to the pericycle). Together, we accurately segmented and analyzed 8849 cells (Table S1).

To compare single cells of a similar developmental state, the cell population of the meristem zone was chosen for analysis (7,859 cells, Fig. S1). To compare between treatments, the mixed-model ANOVA (see methods) was used as it sets a higher benchmark of significance and therefore results in robust and replicable differences between treatments. Before performing the mixed model analysis, the different geometry parameters were transformed to achieve a proxy for normal distribution (Table S2). A corresponding transformation of a given geometry parameter was similarly applied for all tissues (epidermis hair cells, epidermis non-hair cells, cortex, endodermis and pericycle) in all three treatments. The mixed model was fitted for each parameter in each tissue, with treatment and distance from the quiescent center (QC) as a fixed effect and individual plants as random effect. The statistical models were plotted as regression lines for the transformed parameter vs. distance from QC (Fig. 2A-E, Fig S2A-E). This revealed a gradual change in cell volume and surface area, associated with an increase in cell depth and width. Cell length was least affected by distance from QC (Fig. S2F). The data also showed an overall tendency for opposing effects of bri1 and BL treatment on geometric parameters (Fig. 2A-E, Fig. S2A-E). BL treated cells were longer, with reduced radial parameters (i.e. depth and width) while *bri1* cells were shorter, with increased radial parameters. Next, cell geometry was expressed using an anisotropy index (i.e. length^2^/depth*width), where lower values indicate similar cell growth in the longitudinal vs the radial and circumferential axes (isotropic) and higher values indicate increased cell growth in the longitudinal axis (anisotropic). *bri1* cells had the lowest anisotropy and BL cells had the highest, with non-overlapping values between them and a small overlap with WT (Fig. 2F,G, cortical cells are shown). Intriguingly, a parallel analysis of cell volume showed similar values of all treatments (Fig. 2F,G).

**Fig. 2.**
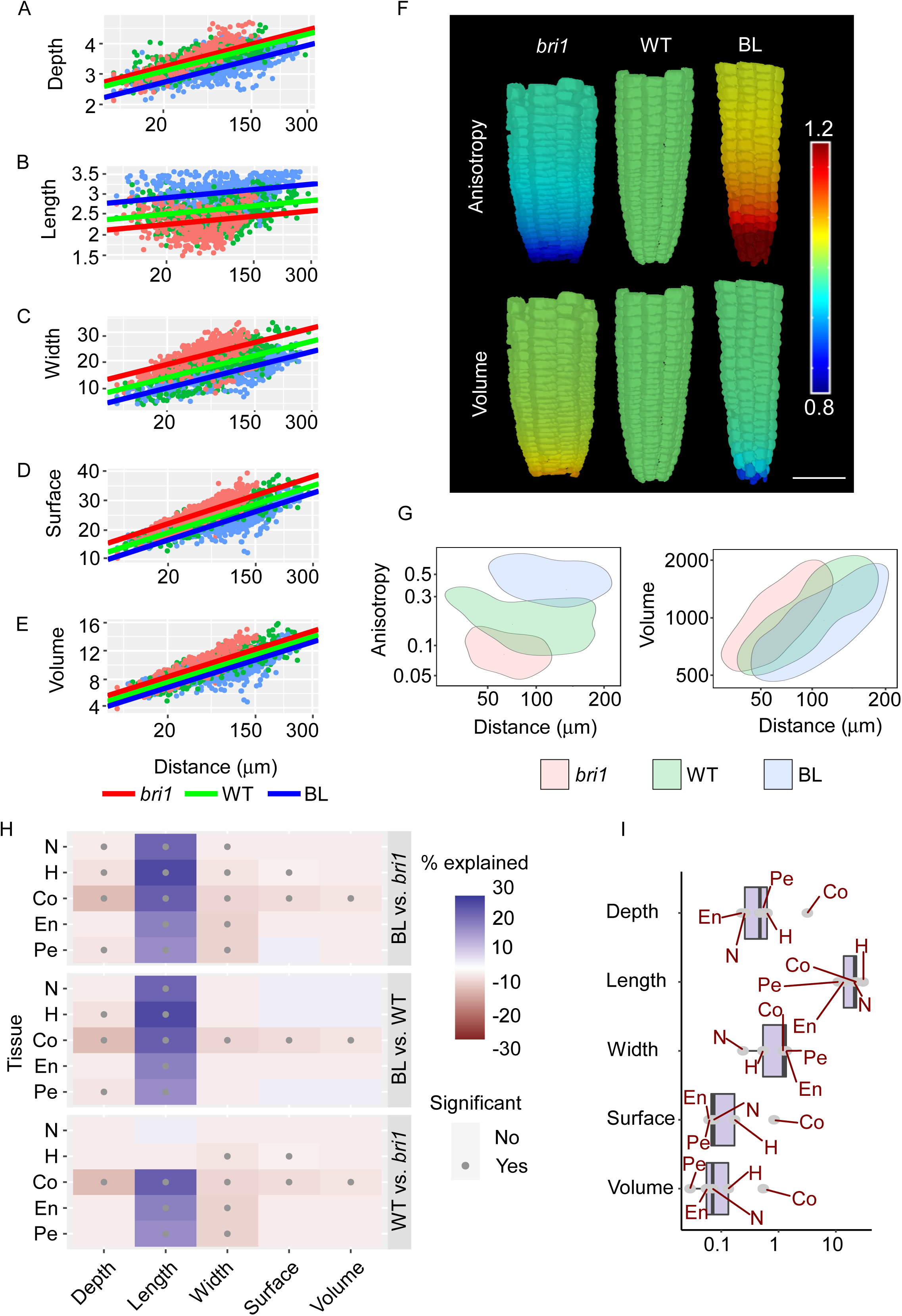
Brassinosteroid signaling has highest effect on cell shape and anisotropy than on cell volume across meristematic tissues (A-E) Robust differences in geometric parameters of individual cells as captured by a mixed model ANOVA (see text for details). Shown here are the effects of the distance from QC and roots with different BR signaling strength (*bri1*, WT and BL-treated WT roots) on cell depth (A), length (B), width (C), surface area (D) and volume (E) in the cortex. Corresponding analyses in other tissues are found in Fig S2. Note the opposite trend between roots with high and low BR signaling. Also note that all geometric parameters, except length are higher in *bri1*. n= WT, 4 roots, 615 cortical cells; *bri1*, 4 roots, 532 cortical cells and BL, 3 roots, 462 cortical cells. (F-G) Comparison of anisotropy index (calculated as length^2^/depth*width, upper roots) and volume (lower roots) between *bri1*, WT and BL-treated WT roots. (F) Display of the cortex tissue in representative segmented roots, depicting relative differences in anisotropy index and volume (WT = 1). (G) Two-dimensional kernel density plots of anisotropy index (left) and volume (right) vs. distance from the quiescent center (QC), of WT, *bri1* and BL-treated WT root cells. Note that BL-treated cells have significantly higher anisotropy, while *bri1* cells have significantly lower anisotropy. In contrast, the two groups of cells showed similar volume values. (H) BR signaling has a higher effect on cell length, depth and width parameters than on cell volume across tissues. Heatmap presenting the percent of variance explained by BR. Shown are all pairwise comparisons organized in three blocks (i.e., WT vs *bri1*, WT vs WT roots treated with BL and *bri1* vs WT roots treated with BL), for 5 geometric parameters in 5 root cell-types and tissues. Blue indicates that the first treatment in the comparison has a higher value. Red indicates that the first treatment in the comparison has a lower value. The higher the opacity, the higher the percent of variance explained. Dot indicates significance (adjusted p-value < 0.05; two-step adaptive correction, see methods). Note the significant, robust, opposing differences between *bri1* and BL-treated samples in length, depth and width across tissues, while cell volume remained largely stable. (I) Boxplot summarizing the effect of treatment on geometric parameters in terms of percent variance explained by BR in a given tissue. N, non-hair cells of the epidermis; H, hair cells of the epidermis; Co, cortex; En, Endodermis; Pe, pericycle.

To determine the magnitude of the BR signaling effect on each geometric parameter, the variance explained by treatment was quantified (Fig. 2H-I, see methods). Notably, when compared to WT, BL treated roots had significantly longer cells in all tissues and reduced cell depth in most of them. By contrast, *bri1* cells were shorter, with significantly higher width in most tissues. When comparing BL to *bri1*, the differences in length, depth and width of cells significantly differed in all tissues, with the exception of endodermal depth, demonstrating a dose-dependent response to BR signaling from *bri1* to WT treated with BL. However, in almost all pairwise comparisons, differences in volume remained non-significant between treatments (Fig. 2H). For simplicity, we quantified and plotted what percent of the variance is the result of treatment (i.e. of BR) for each geometry parameter in a given tissue (Fig. 2I). This demonstrated that volume and surface area were the least affected geometric parameters. Together, this quantitative single-cell geometry analysis demonstrated that BR signaling primarily promotes anisotropic growth. An apparent trade-off between length and depth/width, modulated by the intensity of the BR signaling, ensures cell volume conservation. This trade-off could also be the result of volume serving as a limiting factor.

### 3D time-lapse revealed that the rate and directionality of cellular growth depends on BR and involves a geometry compensation

It remained unclear how these differences in cell shape are generated over time. More specifically, it remained to be determined whether they are a function of the time cells spend in the meristem (duration of growth), the rate of growth in a given axis or a combination of both. As a first step, we performed a kinematic study in 2D to quantify the rate of cell displacement along the root. To this end, the growth of epidermal cells along the meristem and elongation zones was imaged and monitored at 30-min time intervals for a duration of 6h (Fig. 3A, Movie S1, Table S3, Table S4). The analysis revealed a slow displacement of cells (several microns per hour), that gradually increased with distance from the QC. The rate was slower for *bri1* cells (46% slower than WT) and was much faster in the presence of BL (3-fold of WT). Taken together, this direct quantification demonstrated that cells spend more time in the meristem in the absence of BR signaling, and quickly exit it when BR levels are high (Table S5).

**Fig. 3.**
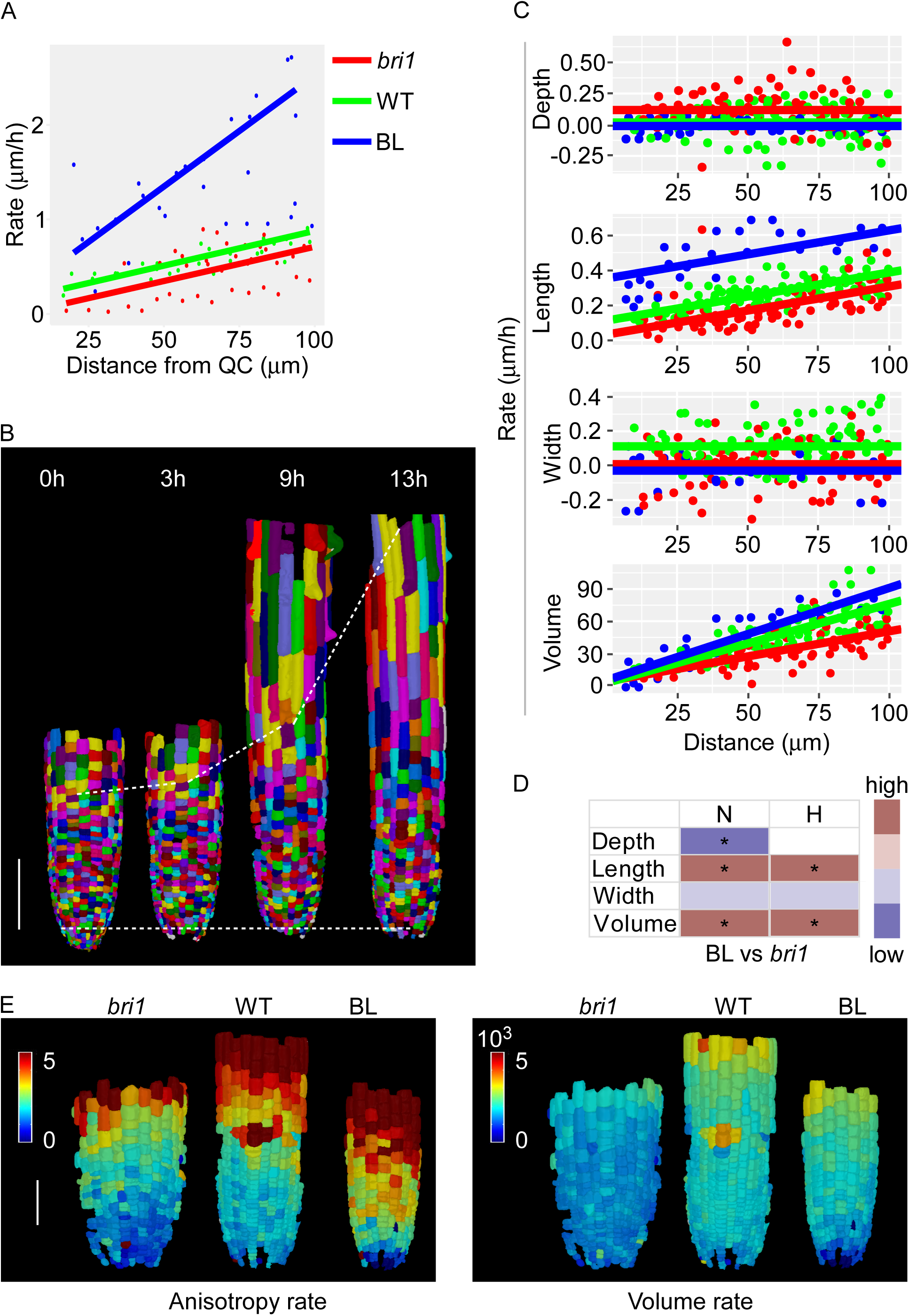
Time-lapse showing that cells with versus without BR have distinct growth rates in alternate directions (A) Rate of epidermal cell displacement along the root meristem. Shown are WT, *bri1* and WT treated with BL. n= WT, 4 roots; *bri1*, 4 roots and BL, 5 roots. (B) 3D segmentation of epidermal cells of WT roots, imaged at 0, 3, 9 and 13 h. Scale bar = 100 μm (C) Single-cell growth in 4D of epidermal (non-hair) cells. Differences in rate of cell growth (depth, length width and volume) were modeled as a function of distance from the QC in WT, *bri1* and WT treated with BL, using analysis of covariance (ANCOVA). n = WT, 130 cells; *bri1*, 119 cells and BL, 68 cells. One root of each treatment. (D) Differences in growth rates upon 4D analysis as in (B) for hair- and non-hair epidermal cells of WT treated with BL and *bri1* samples, summarized as a heatmap. Significant differences are marked by an asterisk. (E) A display of anisotropy rate (left) and volume rate (right) on the corresponding meristematic cells in *bri1*, WT, and WT treated with BL. Scale bar = 50 μm.

Next, we performed 3D time-lapse of WT, WT treated with BL (for 12 h) and *bri1* meristematic epidermal cells at 3h intervals. Epidermal cells were chosen for easier imaging, since image acquisition of the inner tissues under 3D-optimized scanning conditions resulted in a lower signal-to-noise ratio. Due to the slow growth rate of cells in the meristem, the changes made in growth directionality were quantified after approximately 12h (Fig. 3B, Fig. S3, Movie S2, S3, Table S6). As expected, fewer epidermal cells in the longitudinal axis were divided in both *bri1* and BL treated roots as compared to wild type (Fig. S3A-B). No epidermal divisions occurred in other directions. Importantly, while BL dramatically enhanced growth rate in the longitudinal axis, it significantly reduced growth rate in the radial axis, implicated in a lower growth rate of cell width. Consequently, the volume change rate was only slightly higher than WT (Fig. 3C, Fig. S3C). Thus, BL directs longitudinal growth at the expense of radial growth. The growth rate of cell depth was similar to that measured in wild type cells, suggesting that the reduced depth upon BL treatment was primarily the outcome of their short stay in the meristem. In the absence of BRI1, epidermal cells grew significantly more slowly than WT in their longitudinal and width directions and had either higher growth rates than WT in the depth direction (non-hair cells) or the same growth rates as wild type (hair-cells, Fig. 3C, Fig. S3C). Thus, *bri1* cells are larger in depth not only due to the longer time spent in the meristem, but also because of their faster growth rate in this direction. Taken together, meristematic cells with high and low BR signaling have distinct growth rates in opposing directions, which occurs over a distinct time duration (Fig. 3D, Fig. S3D). The overall differences in growth rates and their directionality were implicated in a lower rate of anisotropy growth in *bri1* cells and higher rate after 12h of root exposure to BL, as compared to WT (Fig. 3E). The corresponding cells however had a relative smaller difference than WT. Together, these kinematics and 4D analyses established BR as setting the dynamics of the directionality of growth, where a given axis grows at the expense of the other (Fig. 3C-E, Fig. S3C-D).

### Cell length is inversely correlated with radial growth of the meristem

Moving from the cell to the whole-organ level, we next asked if BL modulation of cell geometry is correlated with radial growth of the meristem. To address this question, radial growth of the different tissues along the meristem was quantified as a function of distance from the QC (as determined by the position of cortical cells along this distance) (Fig. 4A, Fig. S4A-C). While the lateral root cap (LRC) area decreased with distance from QC, all other tissues grew radially, most notably in the stele with higher growth occurring at a distance corresponding to cortical cells 6-8 from QC. This was associated with increased stele and pericycle cell numbers, after which, cell divisions gradually decreased and stopped around the positioning of cortical cell 20 from QC (Fig. S4C).

**Fig. 4.**
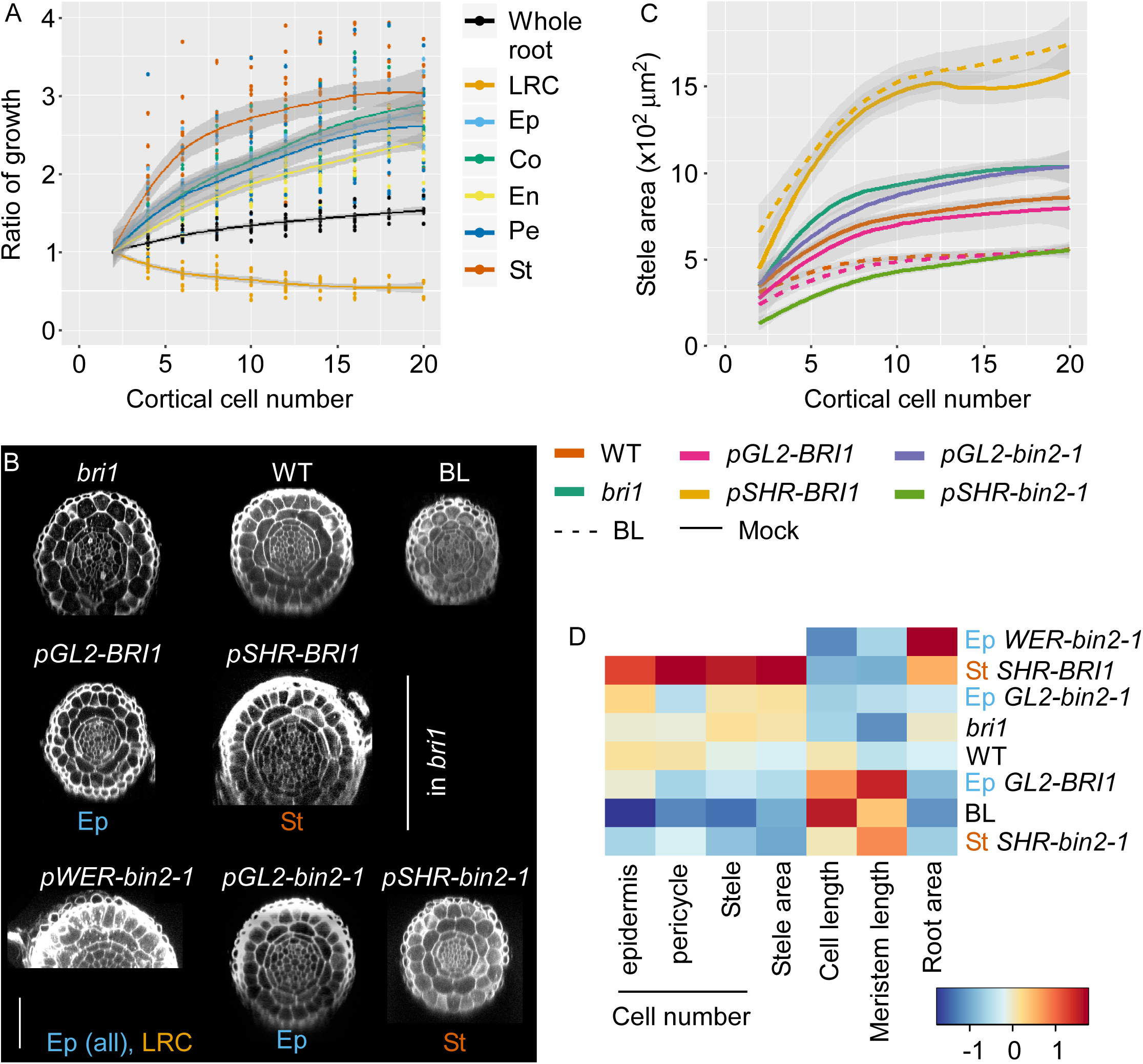
Meristematic cell length and radial size of the meristem are inversely correlated upon BR signaling perturbation. (A) Relative change in radial area of the root meristem and its constituent tissues along the meristem, as indicated by the position of the position of cortical cells in relation to the QC. LRC, lateral root cap; ep, epidermis; c, cortex; en, endodermis; pe, pericycle; st, stele. (n= 24 roots) (B) Confocal images showing radial sections of WT, *bri1* and WT treated with BL, and lines with tissue-specific expression of BRI1 and *bin2-1*, taken 100 μm from the QC. Scale bar = 50 μm. (C) Stele area of lines as in (B), treated or untreated with BL. 3≤n≥11 roots. (D) Heatmap representing mean meristematic parameters of the radial axis (root area at 100 μm, stele area and number of cells in the epidermis, pericycle and the stele) and of the longitudinal axis (average cortical cell length and meristem length) of lines as in (B), treated or untreated with BL. Note the inverse correlation between longitudinal and radial parameters across lines. 5≤n≤11 roots.

Next, we quantified the effect of BR signaling on radial growth and assessed the correlation between this parameter and the BR effect on cell length (as a simple derived parameter of cell geometry) and meristem length. To test if these correlations are affected by mechanical perturbations, lines with tissue-specific perturbation of BR signaling were analyzed (Fig. 4B-C, Fig. S4D-I). These included lines with BRI1 targeted to the epidermis and stele tissues in the *bri1* background using the *pGL2* (directing expression to non-hair cells) and *pSHR* promoters respectively ^16^. WT lines with *bin2-1* (a dominant version of BIN2) driven by the same promoters were used to inhibit BR signaling in these tissues ^19^. As expected, WT treated with BL had a narrow stele, while *bri1* was significantly wider than WT (Fig. 4C-D). Epidermal BRI1 activity was sufficient to limit the stele area of *bri1* and this *pGL2-BRI1* line was not significantly different than WT, in accordance with epidermal control of meristem size ^17 16^. Moreover, radial growth was further restricted in response to BL in this line, similar to WT treated with BL. By contrast, expression of BRI1 in the stele greatly enhanced its size and did not respond to the addition of BL. In contrast to *pSHR-BRI1*, *pSHR-bin2-1* lines had a smaller stele area and *bin2-1* expression in the epidermis (*pGL2-bin2-1*) had a stele area similar to that of *bri1* (Fig. 4B-C, Fig. S4D-I). Roots expressing *bin2-1* under the *pWER* promoter that drives expression in all epidermal cells and LRC (described in ^19^) had a wider meristem than *bri1*, as was seen when BRI1 was limited to the stele (Fig. 3B). Together, BR signaling in the outer and inner tissues had opposing effects on radial growth, negative and positive, respectively. When comparing all lines and treatments, an overall inverse correlation between the size of the radial axis (radial area and cell number) and the longitudinal axis (average cortical meristematic cell length and meristem length) was observed (Fig. 4D). Thus, BR modulation of cell geometry correlates with radial growth of the meristem.

### Simulation model of radial growth in the Arabidopsis root meristem

To understand the mechanism underlying radial growth in the meristem and how it is controlled by BR signaling, we developed a mechanical model using segmented 2D cross-sections. In the model, cell walls are represented as springs connected to vertices that represent the cell junctions (Fig. 5A-C). The walls have a rest length that is initially taken from the starting template, taken at a distance of 8 µm from the QC (Fig. 5A). Turgor pressure is applied to the boundary (it cancels out on inner walls), causing the cell walls to stretch (Fig. 5B). Growth is implemented as a strain relaxation process, with the rest length extension during each growth step implemented as the product of elastic strain times a wall extensibility factor that represents the action of cell wall remodeling gene products (Fig. 5C). In the model, it is possible to individually vary both the elasticity and extensibility factors for each wall, with both affecting growth. Lowering the wall stiffness will cause the wall to stretch more under turgor pressure and increase the growth, as will an increase in the extensibility factor. Although we are modeling the effect of BR signaling on growth, it remains unknown whether BR signaling affects the cell wall stiffness, extensibility factors or a combination of the two ^28, 29^. Here we chose to fit the baseline model to a simple combination of extensibility factors and stiffnesses while matching the WT growth. Note that it is possible to fit the model with a uniform extensibility factor, while only changing the stiffness (or vice versa). To do this, extreme differences in the values for stiffness would be required in different cell layers, as some cells would need to be several hundred-times stiffer than others, likely an unrealistic scenario. From this baseline we model BR signaling changes as differences in stiffness, although other combinations involving differences in extensibility would also be possible. The only cells that divide in the simulations are the cells in the pericycle and stele, which divide at threshold areas based on their cell type (see methods). Since very few tangential divisions occur in the epidermis and endodermis and none in the cortex (Fig. S4D, F), cell division was not modeled for these layers.

**Fig. 5.**
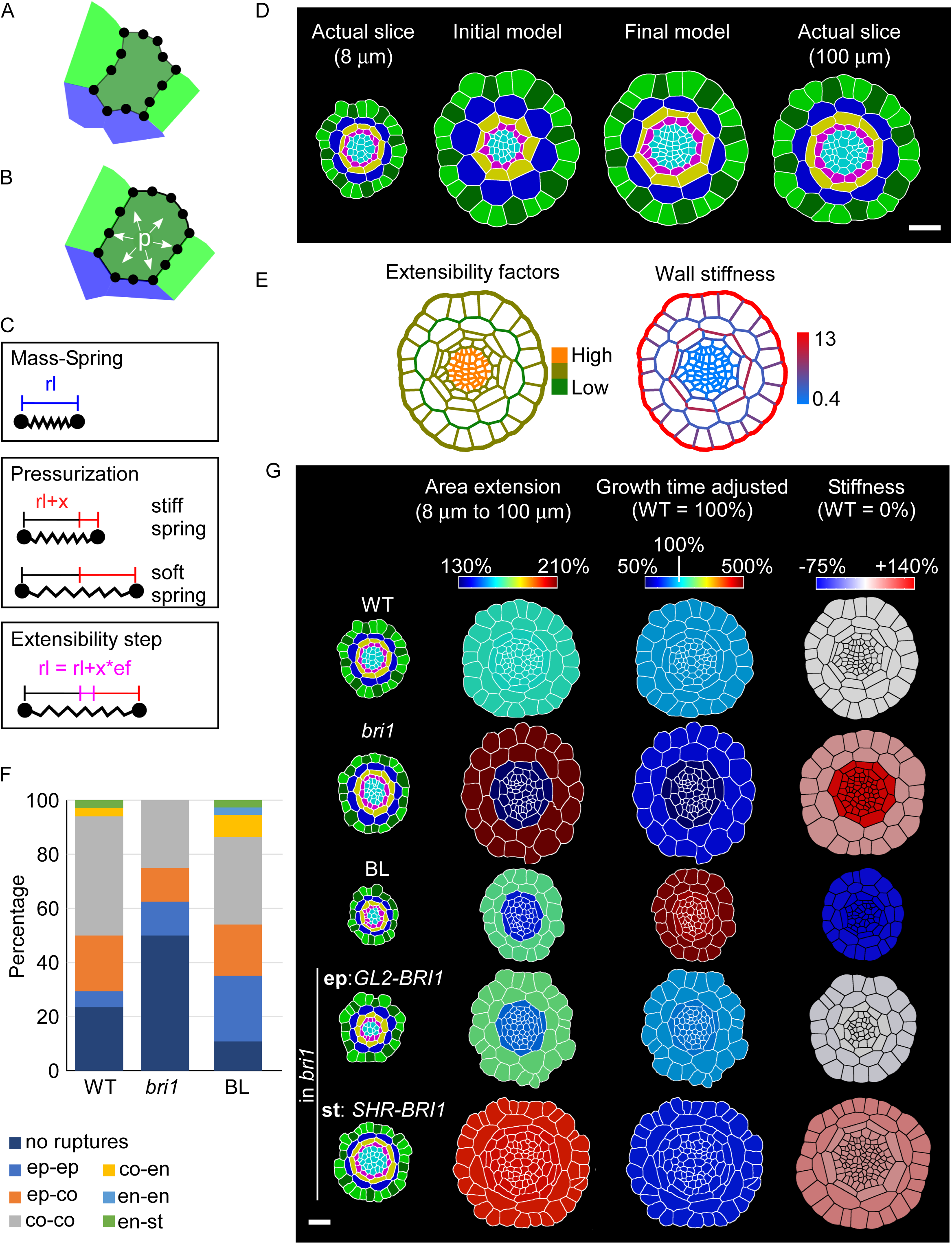
Mechanical model of tissue-specific BR effects on radial growth of the meristem (A) Portion of a segmented mesh used as a model template showing a few cells of the epidermis and the cortex. The cells are colored by cell type: green is epidermis (light green: non-hair cells, dark green: hair cells), blue is cortex. Cell walls are represented by springs that are connected to junction points. A spring has a rest length (rl) defined by the template’s initial geometry. (B) Turgor pressure (p) in the cells puts the springs under tension and leads to elastic expansion (x) of their length. Springs can differ in their stiffness. Here the spring on the outside epidermal wall (in red) is stiffer and expands less than the inner epidermal wall (in orange). (C) Application of extensibility factors (ef) transforms a portion of the expansion (x) into an increase of the spring’s rest length (pink color). (D) Simulation of radial growth requires differential cell wall stiffness. Representative 2D segmentation of a radial optical slice 8 μm from the QC (actual slice 8 μm, left). Simulation model of radial root growth using uniform (second from left, initial model) and differential (second from right, final model) cell wall stiffness. The resulting model output was compared to 2D segmentation of an optical cross-section at 100 μm from the QC (non-virtual slice 100 μm, right). Scale bar = 20 μm. (E) Simulation model for radial growth in the WT root, showing relative distribution of extensibility factors (left) and wall stiffness (right) across cell walls. Note the stiffer outer epidermal and outer endodermal walls, forming the two radial rings. (F) Quantification of cell wall damage (ruptures) in WT, *bri1* and WT treated with BL cells during sample preparation. Note that the epidermis and endodermis are less affected in all samples. *bri1* has the fewest cell wall ruptures while BL-treated roots has the most. (G) Quantification and modeling of the radial growth in WT, *bri1*, BL, BRI1 limited to the epidermis (ep, *pGL2-*BRI1) and stele (st, *pSHR-BRI1*) in the *bri1* background. Cross-sections of the initial templates at 8 μm from the QC, colored by cell type and tissue (left), were compared to their corresponding 2D segmentations at 100 μm and the relative area extension in the outer and inner tissues (second from left, “area extension”). The third column (second from right, “growth time adjusted”) shows incorporation of both time and area extension required for cells to displace from 8 μm to 100 μm (relative to WT). The final column (right column, “stiffness”) shows relative change in stiffness, as compared to WT, in the inner and outer tissues (see Table S9 for numeric data for the heatmap). 3≤n≤8 roots.

The Arabidopsis root meristem reached a relatively stable width at around 15-20 cells from the QC, which corresponds to a distance of 60-80 µm (Fig. 4 and Fig. S4). Thus, we used a cross-section at 8 µm from the QC to represent an early stage of development and another section at 100 µm from the QC to reflect a later stage. This covers approximately 2.57 days (61.86 hours) of WT root growth (calculated from Fig. 3A). The representative cross-section sample was chosen as the closest to the mean value obtained for areal expansion from 8 µm to 100 µm and subsequently used for modeling.

To explore a model with minimal assumptions, we assigned uniform stiffness and extensibility factor to all the cells and ran the growth simulation until the total area matched that of the corresponding 100 µm section (Fig. 5D, Movie S4). Analysis of the tissue-specific areal change found large discrepancies. Cells of the outer tissues (epidermis and cortex) grew much more in the simulation (as compared to the actual, non-virtual 100 µm section) and ended up 12% too large, whereas cells of the inner layers (endodermis and stele) grew less with their final area 31% too small (Fig. 5D, Table S7). This suggests that different tissues must have different stiffness and/or extensibility factors.

It is often thought that the epidermis plays a major role in controlling growth ^30^. Some authors liken plant tissue to a balloon, with the outer epidermal wall restricting growth of the cells within (e.g. ^31 32^). The outer epidermis is also considerably thicker in many plant organs ^30^, including the root meristem ^33^. The flat shape of the endodermal cells also gives the impression that they are constraining the stele. In agreement, among all tissues analyzed the shape of the endodermis was less affected as a function of distance from QC, in particularly cell depth (Fig. S2F-G). In addition, when handling the samples for imaging, occasional ruptures of some of the cells were noted (Fig. 5F).

Quantification of these events revealed that the outer epidermal cell wall fully resisted this damage and that the endodermal walls were less affected in WT, *bri1* and WT treated with BL (Fig. 5F). Interestingly, a larger proportion of *bri1* roots maintained intact cell walls as compared to WT while lower proportion of WT treated with BL maintained intact walls, potentially reflecting distinct cell wall properties controlled by BR signaling. Following these observations, we stiffened the epidermal walls and the outer and inner endodermal walls and lowered the stiffness of the innermost walls in the pericycle and stele (Table S8, Fig. 5D). Finally, we adjusted the extensibility factors to achieve a good fit for the WT sample (Table S8) and achieved a close match (within 0.1%) between the model and the 100µm WT template of the growth of both the inner and outer tissues (Fig 5D, Table S8). Overall, our model can be viewed as a “dual ring” structure, with stiffer epidermal and endodermal walls.

### A plausible model for BR signaling control of radial meristem growth

To model the effects of changes in BR signaling, we selected representative samples of *bri1* and WT treated with BL, as well as lines with BRI1 targeted to the epidermis and stele (i.e. *pGL2-BRI1* and *pSHR-BRI1* respectively). The samples were selected as the closest match to the experimentally observed growth rates of the inner and outer layers. Having established a model for radial growth that involves tissue-specific constraints, we asked to what extent these parameters could reproduce the *bri1* mutant phenotype. Starting with the representative 8 µm *bri1* sample, we grew the template until the total area reached that of the corresponding 100 µm slice. The inner tissue area grew too much (+18%), whereas the outer tissue area grew too little (−6%, Supp. Table S7).

Moreover, the model required 6% more time steps (i.e. relative duration needed for cells to undergo displacement) than the WT to grow to the size of the 100 µm slice, whereas *bri1* requires 76% more time (actual time, Fig 3A) to grow from 8 μm to 100 μm. In addition to *bri1* cells growing slowly along the meristem (Table S7), they could also grow faster in the depth direction (Supp. Fig S2 and Fig. 3). These criteria were met when the stiffness of the cell walls of the inner and outer tissues was increased by 138% and by 42% respectively (Fig. 5G, supp. Table S7). The simulation suggests that BRI1 has a tissue-specific effect on radial growth, the loss of which causes the cell walls of the inner tissues to be considerably more affected than the outer cell walls.

Limiting BRI1 expression to the stele (*pSHR-BRI1*) led to exaggerated radial growth, beyond that of *bri1* (Fig. 4, Fig. S3). Using the mechanical model to explain this non-intuitive phenotypic outcome, we found that the effect on the inner tissues is more prominent than the effect on the outer tissues. A good fit was achieved when stiffness was increased by 32% over WT in the inner walls and by 41% in the outer walls (Table S7, Fig. 5G). In this manner, the extent of *bri1* rescue of mechanical constraint was limited to the inner tissues. This restored the balance of the inner and outer tissues, but all cell walls remained stiffer than in WT. By contrast, when BRI1 was limited to the epidermis (as in *pGL2-BRI1*), the simulation matched the actual growth when the WT stiffness was used for the inner tissues and by softening the outer tissues by 3% (Fig. 5G, supp. Table S7). This is in agreement with the rescue of *bri1* radial and longitudinal parameters (Fig. 4). Indeed, the longitudinal growth rate of *pGL2-BRI1* meristematic cells differed from WT by only 3.7% (Table S7). This suggests that BRI1 signaling in the epidermis is primarily responsible for regulating longitudinal growth ^34^, which has a downstream effect on radial growth in both the inner and outer tissues, to a similar degree.

Finally, we simulated radial growth upon BL treatment while considering that elongation in the meristem is faster (involving compensation by a reduced rate in width) and that the time cells spend in the meristem is shorter than in untreated roots (Fig. 3A, Table S7). Simulation with the WT parameters therefore took too many time steps to reach the final size. This was adjusted by reducing the stiffness of both the inner (by 75.5%) and the outer (by 71.5%, Fig. 5G, Table S7) tissues to match the time and to balance the inner and outer tissue areas. This suggests that BL treatment elicits a similar effect in both the inner and the outer layers, which need to be softened by a similar amount.

## Discussion

Understanding morphogenesis control requires a multiscale analysis of the factors involved. Using the root meristem as the model organ and BR signaling as one of these factors, our work established the key role of BR as controller of cellular growth directionality from the onset of cell production, while cell volume was stabilized. We then linked geometry at the cellular scale to radial meristem growth, and propose a model in which BR signaling controls radial growth via interaction with tissue-specific mechanical constraints.

A longstanding hallmark of BR signaling as concluded from 2D studies, is the promotion of cell elongation in different developmental and physiological contexts ^35^. For example, kinematic analyses of the elongation zone of the root showed that *bri1* cells reach a lower maximal growth rate that ceases early ^16^. Unlike the rapid elongation of cells in the elongation zone, growth of meristematic cells is very slow and thus direct kinematic measurements are limited and require higher spatial resolution (e.g. ^36^). The approach taken here was to generate precise 3D geometry datasets for thousands of meristematic cells across tissues and treatments. These data included time-lapse imaging for quantification of volumetric growth rates in 3D of single cells. This analysis revealed that BR signaling increases cell anisotropy by controlling growth rates in different directions, with a relatively minor effect on cell volume. In response to high BL levels, meristematic cells greatly accelerate their elongation rate while slightly decelerating in width, with the rate of volume increase along the meristem remaining close to the untreated control. When comparing cell shape after long-term exposure to BL, as in our 3D analysis, a lower width becomes significant when comparing to *bri1*. In the absence of BR signaling, growth rates were lower than in WT, with the exception of the depth direction, which was slightly faster, at least in non-hair cells. These findings align with a compensatory process acting on cell geometry, with volume being a primary geometric constraint. Incorporation of time also established *bri1*’s slow rate of cell expansion and associated long cell cycle duration ^13, 16^, also directly demonstrating that these cells spend more time in the meristem and thus have higher total radial growth. Recent studies in the shoot apical meristem suggested that overall stability of meristematic cell volume results from feedback between cell cycle and growth ^37^. However, in the root meristem cell volume gradually increases as cells are displaced from the QC. A computational model proposed that root cells sense their length and stop elongating when reaching a threshold value, depending on BRI1 acting in the meristem ^38^. While several hypotheses are valid, e.g., mechanical strength of the cell which scales with volume and limits cell size ^39^, our data suggest that cells sense a threshold volume and that the target volume increases with distance from the QC. It is plausible that adjusting growth rate in the radial direction, as is achieved by differential BR intensities, is a means of stabilizing a coherent distribution of cell volume. Alternatively, growth directionality can be modulated when volume is a primary geometric constraint.

BR signaling can control directional growth by modulating microtubule arrangement ^40–44^, which in turn guides the positioning of cellulose microfibrils. In the root meristem, a transverse (perpendicular to the root axis) orientation dominates, with the exception of the outer epidermal wall ^45^. The arrangement of the latter becomes transverse in the presence of high BR levels, as in cells leaving the meristem ^44^, suggesting that BR signaling could modulate the anisotropy of the cell wall, which also guides cell shape (e.g., ^46 47, 48^). Earlier experiments using stem segments revealed that BR promotes wall loosening ^28, 29^, in a process involving alteration of its mechanical properties ^28^ and was recently supported in ^49^. When a load was applied to stem segments, higher frequency of breakage was observed in the BL-treated samples, indicating mechanical weakening ^29^. Here, we also observed increased frequency of ruptures when handling root samples treated with BL. In all treatments, these ruptures tended to occur in specific walls, in agreement with differential mechanical properties between tissues. Indeed, differences in growth control among tissues were proposed to be part of root elongation ^50^.

On the organ scale, cell length is affected by tissue-specific BR signaling, with the outer tissues both sufficient and necessary to promote longitudinal cell growth and meristem length, while restricting radial growth. Our simulation model for radial growth in the root meristem provides a step forward towards understanding the mechanistic basis of this outcome, demonstrating that the inner and the outer tissues of the root meristem must have independent regulation of the mechanical parameters of growth, associated with differential radial growth rates. Uniform stiffness and extensibility factors throughout the root meristem are not sufficient to explain the cellular patterning observed *in vivo*. We found that a dual-ring structure, with stiffer epidermal and endodermal cell layers, presents a simple physical arrangement that can regulate growth. Since plant cell growth is thought to be a stress relaxation process, stiffer cell layers would have a dominant role in controlling the growth of the softer layers below them. After fitting the dual-ring model to WT growth rates, we explored how BR signaling can regulate growth in the model. In the *bri1* mutant, the main BR signaling was lost throughout the entire meristem, resulting in a shorter and wider meristem. This phenotype can be interpreted as a trade-off between longitudinal and radial growth, as also observed upon tissue-specific perturbations of BR signaling and as supported by our finding that cell volume is relatively unchanged. However, when considering realistic growth time, it becomes clear that the radial growth is reduced as well (Fig. 5), albeit much less than in the longitudinal direction. We also found that in order to reduce radial growth the walls in the inner tissue had to be stiffened more than those of the outer tissues, meaning that the effect of BR signaling on radial growth is stronger in the inner tissue. Since the outer/inner growth ratio is perturbed in *bri1*, this differential BR effect is required to establish a coherent inter-tissue growth coordination. This also implies that the two rings have a different response to BL. When BRI1 is expressed in the stele only, the meristematic cell length and meristem length are not changed, however, it becomes wider. Again, the initial phenotype seems counter-intuitive, as it opposes the direction of rescue. However, when timing is taking into account in the model, we found that BRI1 expressed only in the stele rescues the balance of the two rings, while all tissues remain stiffer compared to the WT. The largely unaffected longitudinal growth reinforces the notion that BR in the stele mostly affects radial growth. When expressing BRI1 in the epidermis only, both the radial and longitudinal growth are mostly restored in all tissues. Thus, BR affects the inner and outer rings differently, and the outer layer is primarily responsible for longitudinal growth control. Together, our simulations demonstrate how tissue-specific decoding of BR signaling integrates mechanical constraints, thereby shaping the root meristem.

## Methods

### Plant material, growth conditions, and chemical treatments

All Arabidopsis (*Arabidopsis thaliana*) lines were in the Columbia-0 (Col-0) background. The following lines were used: *pGL2-BRI1* and *pSHR-BRI1* ^16^, 35S-eGFP-Lti6b ^51^, *pWER-bin2-1-NeonGreen* and *pSHR-bin2-1-GFP* ^19^, *pGL2*-*bin2-1*-GFP (this study) and *bri1-116*. Seeds were sterilized and germinated on one-half-strength Murashige and Skoog (MS) medium supplemented with 0.2% (w/v) sucrose. Plates with sterilized seeds were stratified in the dark for 2 days, at 4°C, and then transferred to 22°C and to a 16h light/8h dark cycle (70 μmol m^-2^ s^-1^), for 7 days. For chemical and hormone treatments, 3-day-old seedlings were transferred to the relevant supplemented medium and analyzed 4 days thereafter, unless specified otherwise. BRZ and BL were dissolved in 100% dimethyl sulfoxide (DMSO). BRZ and BL were added to a final concentration of 3 mM and 3 nM, respectively. BL activity of the used batch was equivalent to 0.1 nM, corresponding to the activity in our earlier BL batch ^34^.

### Confocal microscopy

For snapshots of live roots, fluorescent signals were detected using a LSM 510 META confocal laser-scanning microscope (Zeiss) with a x25 water immersion objective lens (NA 0.8). Roots were imaged in water supplemented with 10 mg/mL propidium iodide (PI). PI and eGFP were viewed at excitation wavelengths of 488 nm and 980 nm TiSapphier multi-photon, respectively. Fluorescence emission was detected at 575 nm for PI and with a 500 and 530 nm bandpass filter for eGFP. For live imaging (Fig. S2, movie S1), *35S-eGFP-Lti6b* in wild type and *bri1-116* background were imaged using optical plates (Ibidi, 35 mmm-dish) and a LSM 710 inverted confocal laser-scanning microscope (Zeiss) with a x20 air objective lens (NA 0.8). eGFP was viewed with an excitation wavelength of 488 nm and emission detected using a 500 and 530 nm bandpass filter. To prevent roots from drifting, 160 µm channels were molded using acupuncture needles inside MS media supplemented with 2% bacto agar. Seedlings were germinated on 0.5 MS plates and were positioned inside the channels when they were 7-day-old and then flipped over to fit inside the optical plates. In case of chemical or hormonal treatment, the treatment was added to the bacto agar-supplemented MS during preparation.

For 3D segmentation, roots were fixed using the mPS-PI protocol ^52^, and placed in a chamber made of 200 μm-thick dual-sided tape (to prevent the sample from changing shape due to cover slip pressure). The tape was glued to a slide, roots were placed inside and a cover slip was glued above. Imaging was conducted using a LSM 510 META confocal laser-scanning microscope with a x40 oil immersion objective lens (NA 1.3). PI was viewed with an excitation wavelength of 488 nm and emission collected with a 575 nm bandpass filter, using a fine pinhole of 1 nm and Z-steps of 0.8 nM. For 4D image acquisition, molds with channels as described above were made to fit a microscope slide. Roots of 7-day-old seedlings were positioned in the channels, closed with a cover slip and kept vertically in the growth chamber between imaging sessions. Imaging was performed every 3 hours.

### Segmentation analysis

All acquired images underwent pre-segmentation processing, including background subtraction and signal enhancement, using Fiji. MorphoGraphX ^27^ was used for all segmentations (i.e. 2D, 3D and 4D) followed by manual corrections. Images were segmented using the Insight Toolkit (ITK) morphological watershed processes in MorphoGraphX and a volumetric (3D) mesh was extracted. 3D meshes were analyzed using the 3D Cell Atlas pipeline ^53^, which uses an organ centric coordinate system to assign directions to each cell. Cell sizes (length, width and depth) were obtained by measuring the size of each cell along these directions through the cell’s center of gravity. Cell geometry data were exported to csv files and further analyzed in R. For 4D analysis, lineage tracking was used to track cells between time points. Cell divisions were extracted based on lineage assignment. 4D movies were generated using Abrosoft FantaMorph software using the first and the last time points segmentations, and lineage representation.

### Radial analysis

Fixed roots were imaged with the microscope settings used for the 3D analysis and loaded into Fiji. Roots were straightened and a horizontal line was drawn at the point of the intended slicing. The dynamic reslice Fiji tool was used to generate an optical cross-section.

### Classification of meristematic cells

Cells were classified into meristem and elongation zone cells as follows. Using the expectation maximization (EM) algorithm as implemented in the mixtools R package ^54^, a two-Gaussian mixture model was fitted to the cell length parameter in each combination of tissue and BR condition (WT and each of the BR perturbations) in roots. The probability to be in the short-length Gaussian was calculated for each cell. Cells with probability >0.8 were considered meristem cells. Further analyses were performed on the meristematic cells.

### Statistical analysis and quantification of BR contribution to the variance

For 3D data, the experimental design had fixed (i.e., BR perturbations and distance from the QC) and random (i.e., biological replicate) factors. Therefore, hypothesis testing was performed with the mixed model ANOVA using the R lmer function ^55^. ANOVA assumes normal distribution. Hence, we visualized the data and if required, transformed parameters to achieve a proxy for normal distributions. The transformations are detailed in Table S1. Post-hoc tests were performed using the Tukey procedure, as implemented in the multcomp package ^56^, in each model that was significant after correction for multiple hypotheses. Correction for multiple hypotheses was performed using a two-step adaptive procedure ^57^. Briefly, in the first stage we used the BH procedure with α=0.05.

This stage resulted in 17 significant models out of the 25 possible models. In the second stage, we performed the BH procedure on the post-hoc comparisons with α =0.05*17/25. This procedure guarantees that the false discovery rate (FDR) is kept at 0.05 throughout the entire experiment. To determine the percentage of variation explained by BR, we first used the Insight package ^58^ to calculate the variance explained by the fixed factors, the random factors and the residual variance. Then, to isolate the variance explained by BR conditions we multiplied the fixed variance by the proportion of the sum of squares of the BR conditions.

The 2D data were analyzed with using a mixed model ANOVA, as described for the 3D data. Since we were interested only in a small part of all possible genotype comparisons, we generated a model for each pair of interest. As a result, no post-hoc test was required.

For the 4D data, each cell was measured at two time points. We selected cells located up to 60 μm from the QC at the first time point and up to 100 μm in the second time point. This ensured that cells in the comparison were residing in the meristem throughout the experiment. Analysis of covariance (ANCOVA) was performed using the “lm” function ^59^ for each geometrical parameter, with treatment and distance-from-QC as main factors. Each ANCOVA was started with an interaction model, which was then simplified when possible, as described in the R book ^60^. Post-hoc comparisons between treatments were performed using the contrast package ^61^.

### Kinematic analysis

Roots were positioned in channels on an optical plate (as previously described) and imaged at 30-minute intervals, over 6 hours. The resulting images underwent stitching and regression correction to overcome 3D drifts of the sample (all within Fiji). The same cell was traced and its distance from QC at t=0 and t=6h was recorded. Measurements were plotted and a slope was derived from the fitted curve.

To calculate meristem dynamics in WT, *bri1, pGL2-BRI1* and *pSHR-BRI1*, cell production rate was evaluated on day 6-7. This was performed by dividing the length of root elongation between days 6 and 7 by the average mature cell length. Next, we measured average meristematic cell length and used this value to calculate the total “length” that left the meristem in this time period (i.e. average meristematic cell length * cell production rate). Using the meristematic cell number, we calculated meristem length (meristem cell number * average meristem length) and used it to calculate growth rate (meristem length / length that left the meristem). A good match was found between the calculated ratio of WT vs *bri1* and the ratio derived directly from live imaging (Table S4). Therefore, we applied the calculated method for the genotypes that were not subjected to a direct kinematics analysis.

### Model methods

Model simulations were done on radial cross-sections at 8μm and 100μm distances to the QC, that were extracted from 3D images using Fiji. The 2D cross-section images were loaded into MorphoGraphX and segmented, starting with a square mesh in the XY plane covering the entire cross-section. The signal from the 2D cross-section was projected onto the mesh with a triangle size slightly smaller than the pixel size of the original image. The meshes were then manually seeded and segmented using the watershed segmentation process. In a final step, the mesh walls were smoothed. For the different genotypes and treatments (WT, *bri1*, BL*, pSHR-BRI1*, and *pGL2-BRI1*) we segmented between 3 and 8 replicates, from which we chose one representative sample for each genotype based on their sum of relative mean square errors of the growth of the outer (epidermis, cortex) and inner (endodermis, pericycle, stele) tissues. The samples with the smallest error scores were selected as representative samples, with the exception of the *bri1* sample, where we chose the second-best sample, as the best sample showed an overall asymmetry in the epidermis due to fewer LRC layers in one side of the root at 8 μm. The mass-spring model was implemented using MorphoDynamX (www.MorphoDynamX.org), a modeling platform based on MorphoGraphX. Models were stored in VLab ^27, 62^. The main parameters of the mass-spring model were spring stiffness, extensibility factors (both assigned to the wall edges, see Fig 5A) and max cell area (assigned to cells). See Table S8 for an overview of the stiffness and extensibility factors of the model and Table S7 for changes required to model the treatments and genotypes. The maximum cell area of stele was 30 µm^2^ and was 70 µm^2^ for the pericycle.

## Acknowledgments

We thank O. Hamant, T. Shemesh and I. Efroni for their comments on the manuscript. We thank A. Farhat for excellent technical assistance, the Life Sciences and Engineering Infrastructure Center (N. Dahan, Y) and the Russell Barrie Nanotechnology Institute at the Technion. This research was supported by grants from the Israel Science Foundation (1725/18) to SS-G, core funding from the Max Planck Institute for Plant Breeding Research to RSS and a BBSRC Institute Strategic Programme GEN [BB/P013511/1] grant to the John Innes Centre.

## Supplementary Figure Legends

**Fig. S1.**
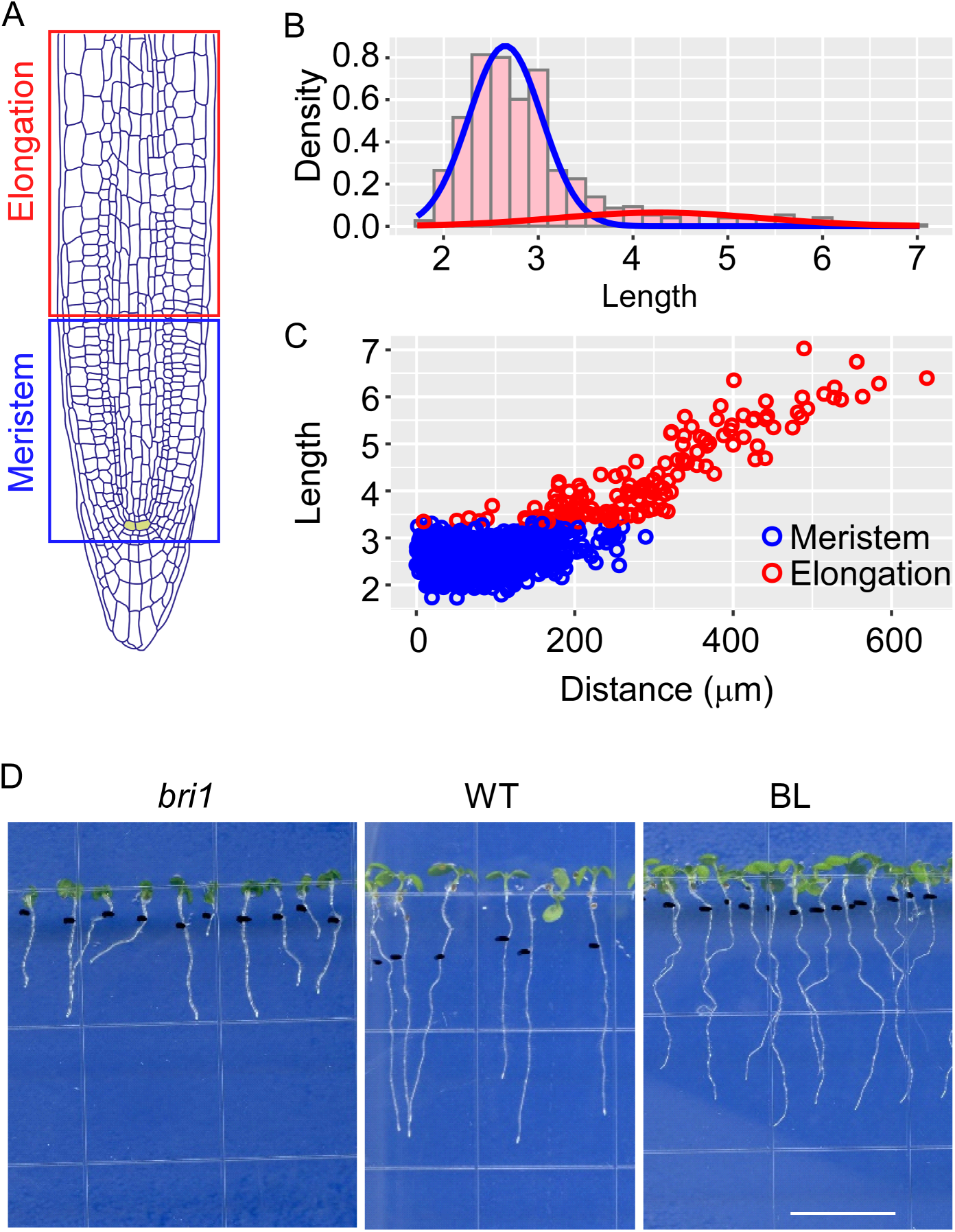
Algorithm-based labeling of meristematic cells and elongating cells (for quantification of single-cell geometry). (A) Schematic presentation of the meristem and elongation zone. The quiescent center (QC) cells are marked in yellow. (B) Gaussian mixture model of cell length captures two populations of relatively short (blue gaussian) and long (red gaussian) cells. (C) Scatter plot of the cells in the upper panel, depicting cell length vs. distance from the QC. Each dot represents a single cell of the meristem (blue) and elongation (red) zone. An example is shown for cells of WT cortex tissue. (D) Morphology of 7-day-old seedlings of *bri1*, WT and WT treated with BL for 4 days. Scale bar =10 mm

**Fig. S2.**
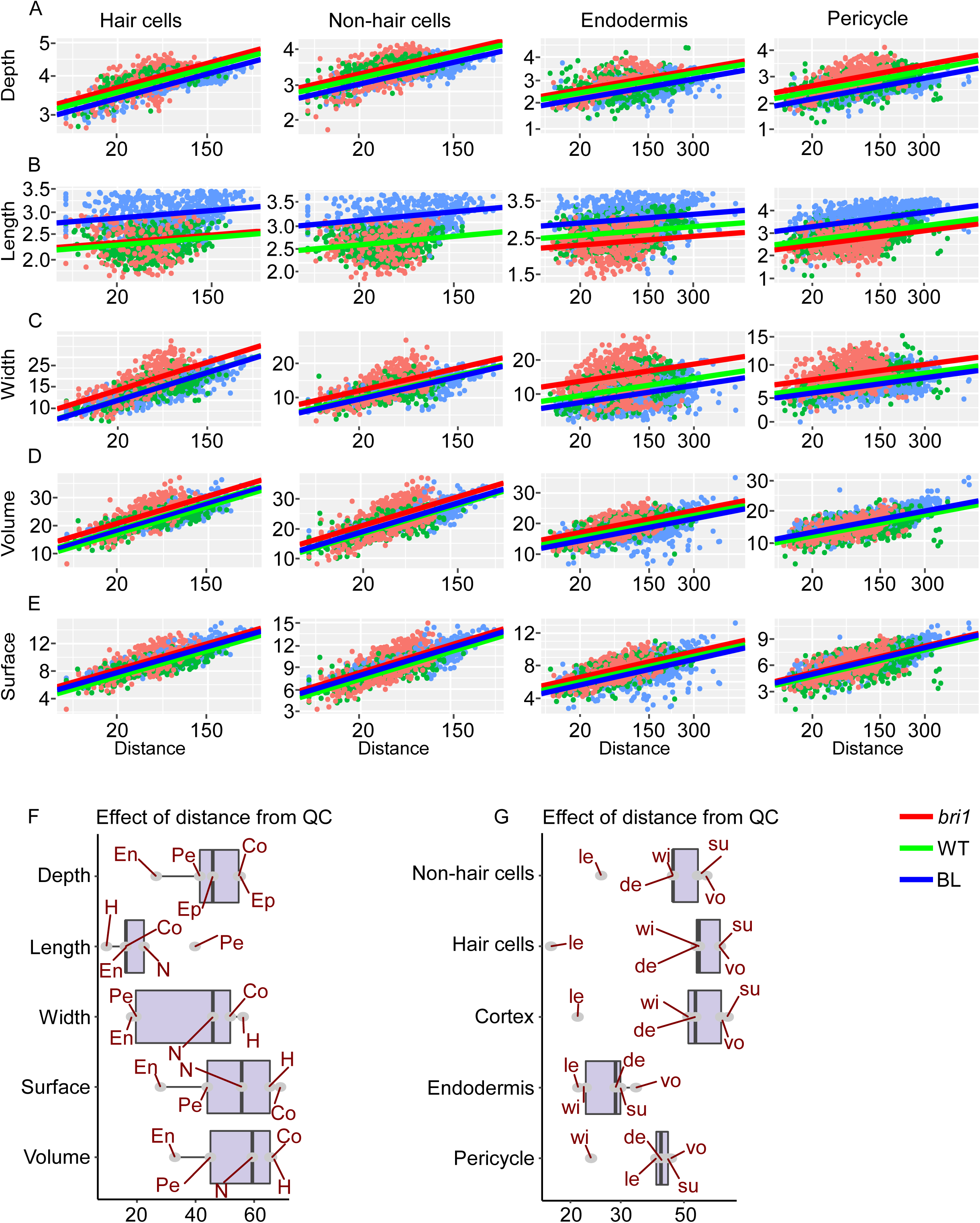
Low and high BR signaling show opposing trends in geometry parameters across meristematic tissues (A-E) Differences in single-cell geometric parameters. Shown are the effects of the distance from QC and roots with distinct BR signaling strength (i.e., WT, *bri1* and BL treated roots) on length (A), depth (B), width (C), volume (D) and surface area (E) in epidermis (hair- and non-hair cells), endodermis and pericycle cells. Note the opposite trend between roots with high and low BR activity. (n epidermis, N cells: WT – 374 cells; *bri1* – 376 cells; BL – 280) (n epidermis, H cells: WT – 288 cells; *bri1* – 322 cells; BL – 460 cells) (n endodermis: WT – 445 cells; *bri1* – 389 cells; BL – 622 cells) (n pericycle: WT – 633 cells; *bri1* – 749 cells; BL – 833 cells). (F,G) Boxplots summarizing the percent variance explained by distance from QC, for each geometric parameter in a given tissue. The percent of variance is estimated from R2. Boxes are grouped by geometric parameter (F) or by tissue (G). Note that length is the geometric parameter and endodermis is the tissue with the least dependence on distance.

**Fig. S3.**
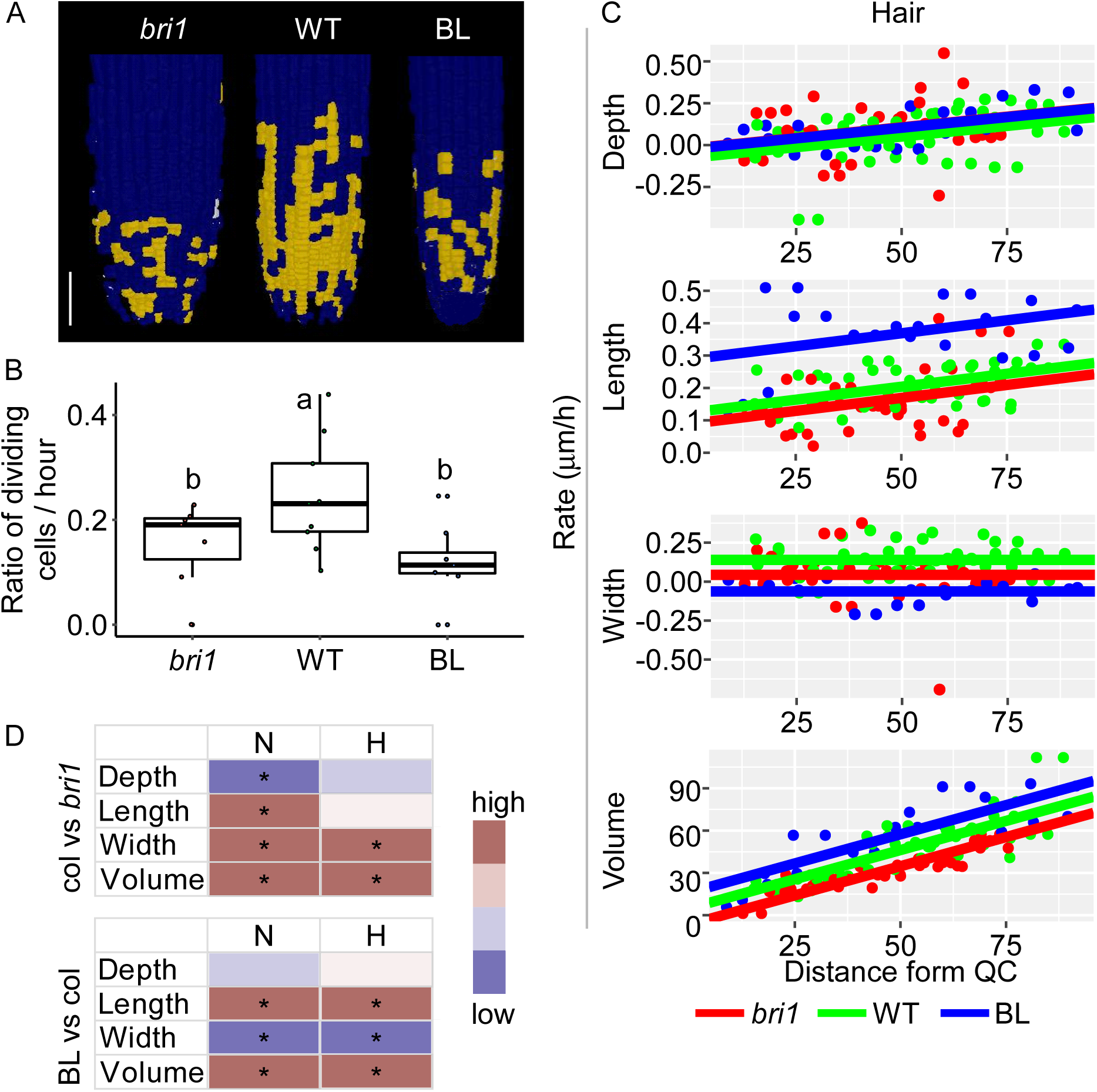
Cell division and growth of individual cells in 3D over time (A) Epidermal cells that underwent divisions in *bri1*, WT and WT treated with BL after 10.5h, 13h and 12h of imaging, respectively, are marked in yellow, scale bar = 50 μm. (B) The fraction of meristematic cells that underwent divisions is shown as the number of mother cells divided by total meristematic cells, normalized to imaging duration. (C) Differences in expansion rate (depth, length, width and volume) of hair-cells were plotted as a function of distance from the QC in WT, *bri1* and WT treated with BL, using analysis of covariance (ANCOVA). n = WT, 76 cells; *bri1*, 68 cells and BL, 96 cells of 1 root. (D-E) 4D analysis of differences in growth rates of (C) WT and *bri1* hair- and non-hair epidermal cells, (D) WT cells treated with BL and (E) WT cells, summarized as a heatmap. Significant differences are marked by an asterisk.

**Fig. S4.**
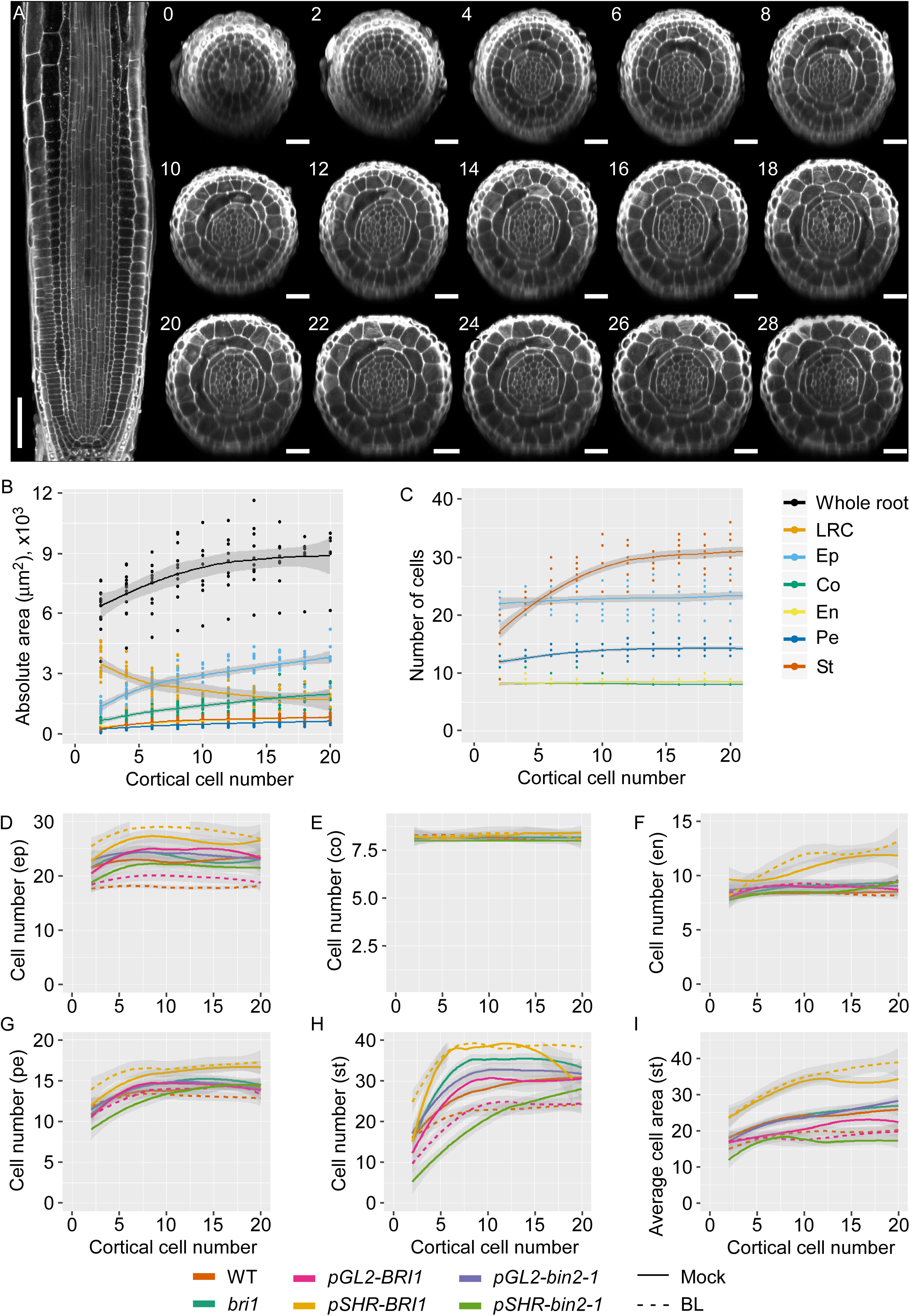
Quantification of growth parameters in the radial axis and their modulation by BR (A) Confocal images of a WT root and corresponding sequential radial section taken at the QC, marked by the position of the cortical cell (2 closest to the QC). Longitudinal scale bar = 50 μm, radial scale bar = 20 μm. (B) Radial WT root area, as measured from radial cross-sections of the different tissues composing the root at a given distance from the QC, marked by position of cortical cells. n= 24 roots. (C) Number of cells in radial cross-sections of the different tissues composing the WT root, at a given distance from the QC, marked by position of cortical cells. (D-H) Number of cells in radial cross-sections of the different tissues composing the root, at a given distance from the QC, marked by position of cortical cells in WT, *bri1* and WT treated with BL, and lines with tissue-specific expression of BRI1 and *bin2-1*, treated or untreated with BL. (I) Average area of stele cells of lines as in D-H. 3≤n≤11 roots LRC, lateral root cap; ep, epidermis; co, cortex; en, endodermis; pe, pericycle; st, stele

## Supplemental Movies

**Movie S1** Growth of WT, *bri1* and BL treated roots

Root growth of WT, *bri1* and BL treated roots, harboring 35S-Lti6b-eGFP. Roots were imaged every 30 minutes for a total duration of 6 hours. Movies were acquired using optical channels under inverted confocal microscope. STD Z-projection is presented. These movies were used to measure meristematic cell displacement (Fig. 3A).

**Movie S2** 3D-segmentation over time

WT, *bri1* and BL treated roots were imaged for 13, 10.5 and 12 hours, respectively. Images were segmented and initial and final images were morphed to represent root growth (see also Fig. 3 and S3). Elongation and meristem zones are shown.

**Movie S3** 3D-segmentation of meristematic cells over time

Movie as in S2 showing meristematic zone. Note the faster progression of meristematic cells upon BL treatment.

**Movie S4** Model of WT radial growth using equal wall stiffness

Representative WT root was used to model radial growth from section located at 8 μm from QC to 100 μm from QC. A uniform stiffness and extensibility factor were assigned to all the cells, resulting in deviation from the actual shape of the cross section (see Fig 5D, Table S7).

**Movie S5** Simulation of BR signaling control of radial meristem growth

Radial growth simulation of WT, *bri1*, BL treated roots, *pGL2-BRI1* and *pSHR-BRI1* (See Fig. 5).

**Table S3.** Kinetics table

| Line | Kinetics | Vs. WT |
| --- | --- | --- |
| WT | 0.0077 | 1.00 |
| *bri1* | 0.0044 | 0.569 |
| BL | 0.023 | 3.00 |

**Table S4.**
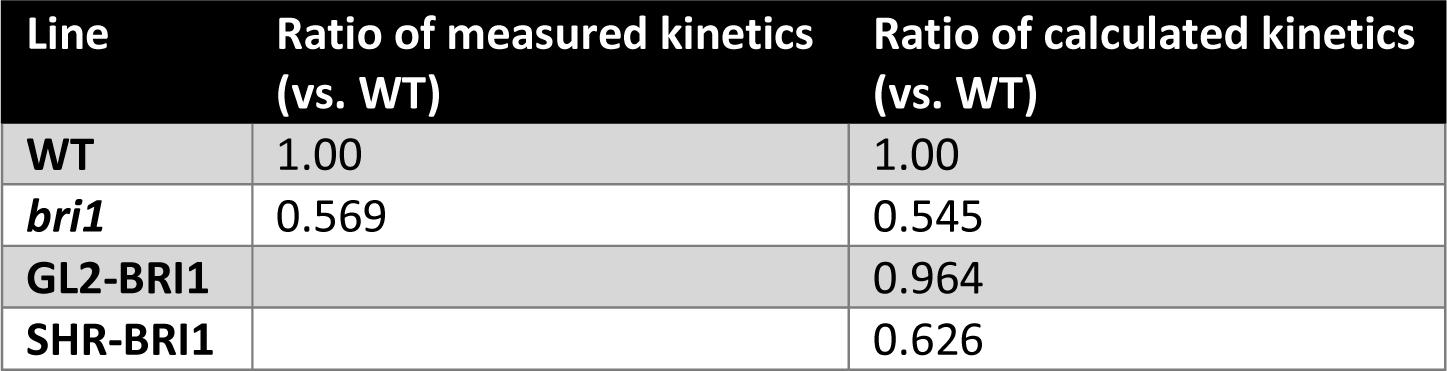
Calculated kinetics table

**Table S5.** meristem duration

| Line | Meristem length ($\mu\text{m}$) | Rate ($\mu\text{m/h}$) | Duration (hours) | Duration (days) |
| --- | --- | --- | --- | --- |
| WT | 194 | 1.49 | 130.72 | 5.45 |
| *bri1* | 112 | 0.49 | 229.89 | 9.58 |
| BL | 234 | 5.83 | 40.16 | 1.67 |

**Table S7.** simulation parameters for all lines

| Model | Target time to WT | Parameters vs. WT | Parameters used | Stiffness to WT |  | Stiffness to *bri1* |  | Time steps | To WT time | To target time | Inner ratio | Outer Ratio | I/O |
| --- | --- | --- | --- | --- | --- | --- | --- | --- | --- | --- | --- | --- | --- |
|  |  |  |  | Ratio Inner | Ratio Outer | Ratio Inner | Ratio Outer |  |  |  |  |  |  |
| WT | 0% | same | same growth rate & stiffness | - | - | - | - | 1861 | - | - | 0.69 | 1.12 | -38.73% |
|  |  | I 0 ; O 0 | WT | 0% | 0% | -55% | -30% | 6847 | 0.0% | 0.0% | 0.999 | 1.000 | -0.001 |
| bri1 | 75.7% | I 0 ; O 0 | WT | 0% | 0% | -58% | -30% | 7227 | 5.5% | -39.9% | 1.176 | 0.940 | 0.252 |
|  |  | I+138 ; O+42 | bri1 | 138% | 42% | 0% | 0% | 12032 | 75.7% | 0.0% | 0.999 | 1.000 | -0.002 |
| BL | -78.9% | I 0 ; O 0 | WT | 0% | 0% | -58% | -30% | 8781 | 28.2% | 506.9% | 1.057 | 0.979 | 0.080 |
|  |  | I-75.5 ; O-71.5 | BL | -75.5% | -71.5% | -90% | -80% | 1450 | -78.8% | 0.2% | 1.000 | 1.000 | -0.001 |
| GL2 | 3.7% | I 0 ; O 0 | WT | 0% | 0% | -58% | -30% | 7293 | 6.5% | 2.7% | 1.014 | 0.996 | 0.019 |
|  |  | I+138 ; O+42 | bri1 | 138% | 42% | 0% | 0% | 12040 | 75.8% | 69.5% | 0.838 | 1.048 | -20.0% |
|  |  | I 0 ; O-3 | GL2 | 0% | -3% | -58% | -32% | 7076 | 3.3% | -0.4% | 1.000 | 1.000 | 0.000 |
| SHR | 11.3% | I 0 ; O 0 | WT | 0% | 0% | -58% | -30% | 7611 | 11.2% | -30.4% | 0.955 | 1.024 | -0.067 |
|  |  | I+138 ; O+42 | bri1 | 138% | 42% | 0% | 0% | 12770 | 86.5% | 16.8% | 0.816 | 1.098 | -25.7% |
|  |  | I+32 ; O+41 | SHR | 32% | 41% | -45% | -1% | 11020 | 60.9% | 0.8% | 0.997 | 1.001 | -0.004 |

**Table S8.** simulation parameters for WT

|  |  | Outer |  | Inner |  |  |
| --- | --- | --- | --- | --- | --- | --- |
| ***Stiffness*** |  | **1** | **2** | **3** | **4** | **5** |
| **Outside** | **0** | 13 | - | - | - | - |
| **Epidermis** | **1** | 6.1 | 3.7 | - | - | - |
| **Cortex** | **2** | - | 3.5 | 9 | - | - |
| **Endodermis** | **3** | - | - | 2.5 | 3.9 | - |
| **Pericycle** | **4** | - | - | - | 1.1 | 1.1 |
| **Stele** | **5** | - | - | - | - | 0.4 |

| ***Growth Factors*** |  | **1** | **2** | **3** | **4** | **5** |
| --- | --- | --- | --- | --- | --- | --- |
| **Outside** | **0** | 0.001 | - | - | - | - |
| **Epidermis** | **1** | 0.001 | 0.0002 | - | - | - |
| **Cortex** | **2** | - | 0.0002 | 0.001 | - | - |
| **Endodermis** | **3** | - | - | 0.001 | 0.001 | - |
| **Pericycle** | **4** | - | - | - | 0.005 | 0.005 |
| **Stele** | **5** | - | - | - | - | 0.005 |

**Table S9.** Area Extension

|  | Area Extension (Ratio) |  | Area Extension (%) |  | Time Factor | Area Extension (Time Adjusted) |  |
| --- | --- | --- | --- | --- | --- | --- | --- |
|  | Outer | Inner | Outer | Inner |  | Outer | Inner |
| WT | 2.63 | 2.62 | 163.29 | 161.92 | 1.00 | 2.63 | 2.62 |
| *bri1* | 3.13 | 2.28 | 213.12 | 128.26 | 1.76 | 1.78 | 1.30 |
| BL | 2.68 | 2.42 | 167.56 | 142.47 | 0.21 | 12.66 | 11.47 |
| *pGL2-BRI1* | 2.69 | 2.49 | 168.94 | 149.32 | 1.04 | 2.59 | 2.40 |
| *pSHR-BRI1* | 2.98 | 2.99 | 197.61 | 198.97 | 1.60 | 1.86 | 1.87 |

## Notes

### Competing Interest Statement

The authors have declared no competing interest.

